# Identification of PTBP1 responsible for caspase dependent YRNA cleavage

**DOI:** 10.1101/298851

**Authors:** Jun Ogata, Yuki Sugiura, Akinori Kanai, Masafumi Tanaka, Hirotaka Matsui, Masato Ohtsuka, Toshiya Inaba, Motoyuki Otsuka, Ai Kotani

## Abstract

Some RNAs such as 28S rRNA, U1 snRNA, and Y RNAs are known to be cleaved during apoptosis. As the underlying mechanism is yet unclear, the functions and biological significance of RNA degradation in apoptosis remain elusive. We previously identified novel, functional small RNAs named AGO-taxis small RNA (ASR) that are specifically bound to AGO1. Here, we investigated ASR biogenesis, which appears to be non-canonical. Y RNAs, non-coding RNAs degraded during apoptosis, were identified as the precursors of several ASRs. Cell-free analysis combined with fractionation methods revealed that the apoptosis-specific biogenesis of ASRs or Y RNA degradation was induced by PTBP1—an endoribonuclease inhibitor of Y RNAs. PTBP1, a splicing factor, was truncated by caspase 3, which subsequently activated endoribonuclease to induce biogenesis of ASRs and Y RNA cleavage.

---

Apoptosis is essential for the development and homeostasis of metazoans (1). In an apoptotic cell, some components are decomposed by its own proteases or nucleases (2,3). While the processes associated with chromatin fragmentation, cytoskeleton disruption, nucleus condensation, and membrane blebbing are extensively studied and well understood, the mechanisms and functions of apoptotic RNA degradation remain unelucidated (4,5).

Active RNA cleavage in apoptotic cells is observed for a few non-coding RNA species such as 28S rRNA, U1 snRNA, and Y RNAs. 28S rRNA, a part of the large subunit of the ribosome, is transcribed by RNA polymerase I in the nucleolus. While 18S rRNA remains intact during apoptosis, 28S rRNA is immediately fragmented (6). U1 snRNA, RNA forms U1 small nuclear ribonucleoprotein (U1 snRNP) complex is involved in mRNA splicing, is also cleaved in apoptotic cells (7). During apoptosis, one of the components, U1-70K, is cleaved by the activated caspase 3 and the single-stranded 5′-end is simultaneously truncated by an unknown ribonuclease (8). In addition, Y RNAs undergo rapid cleavage by caspase 3 during apoptosis (9). Y RNAs are non-coding RNAs that are widely conserved from bacteria to mammals (10). Humans have four Y RNAs: Y1, Y3, Y4, and Y5 RNA. In the nucleus, Y RNAs contribute to DNA replication initiation through formation of the replication origin (11,12). Y RNAs form Ro-RNP complex with Ro60, which has been reported to be one of the common autoantigens observed in autoimmune diseases, to regulate the cytoplasmic localization of Ro protein by masking the nuclear localization signal of the protein (13). Although these cleavage activities were observed more than 20 years ago, the underlying mechanisms and identification of the responsible ribonuclease have remained elusive. Autoantibodies against these ribonucleocomplexes (ribosome, U1 snRNP, and Ro-RNP) are often detected in the sera of patients with systemic autoimmune diseases, indicative of the involvement of apoptotic RNA cleavage in these pathological conditions (14–17).

The study of small RNA, including microRNA (miRNA), endosiRNA, and piRNA, has greatly expanded after the emergence of next-generation sequencing (18–20). We previously identified a new class of small RNAs that specifically bind to AGO1, but not AGO2, prepared from Epstein-Barr virus (EBV)-positive lymphoma cells, by small RNA sequencing (21). These small RNAs named AGO-taxis small RNAs (ASRs) exhibited RNA interference activity and dramatically increased during the EBV lytic infection, which was accompanied with massive cell death.

To investigate the correlation and biological significance of this small RNA cleavage, apoptosis, and pathogenesis of autoimmune diseases, it is crucial to elucidate the detailed mechanism of RNA cleavage in apoptosis.

Here, we conducted a cell-free assay in combination with some fractionation methods to reveal that Y RNA cleavage in apoptosis is caused by a ribonuclease and its inhibitor PTBP1—a well-known splicing factor (22). PTBP1 induced apoptosis-dependent Y RNA cleavage through its truncation by activated caspase 3. Following PTBP1 cleavage, Y RNA single-strand region may be exposed to any kind of endoribonucleases.

## Results

### Agotaxis small RNAs are derived from the stem regions of Y RNAs

The consensus motif of ASRs was already identified (5) and UUGACU, known to be conserved in the stem region of the non-coding Y RNA, was identified as the binding motif of Ro autoantigen. A common structure of Y RNA was drawn using secondary structure of Y RNAs (23). The motif was plotted on the common secondary structure (Figure 1A) and small RNAs were found to be derived from the 3′ stem regions of Y RNAs via cleavage of the single-strand loop region.

**Figure 1.**
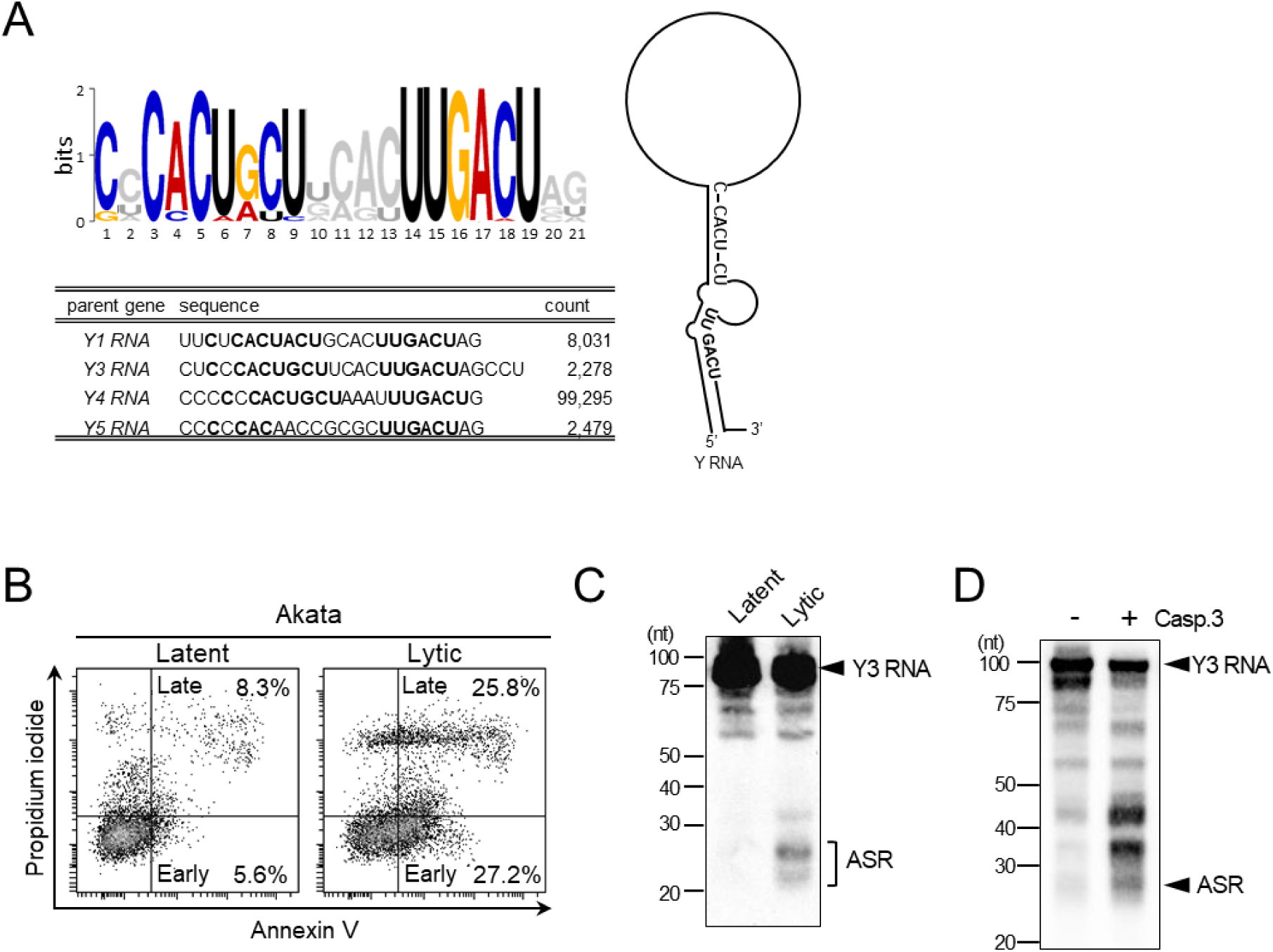
Y RNA cleavage is activated in apoptotic cells. (A) Common motif of ASRs was identified by MEME motif search. The motif, indicated in the capital letter, was plotted on Y RNA common structure. Sequences of the most abundant ASRs from each Y RNA were analyzed. (B) EBV lytic infection was induced by BCR stimulation using anti-human IgG antibody. After 24 h, apoptosis induction was investigated by flow cytometry. Annexin V single positivity shows early apoptotic cells and double positivity shows late apoptotic cells. (C) Y3 RNA and ASR cleavage product were detected by northern blotting. An arrow head shows the full-length Y3 RNA; bracket, ASR. (D) The cytoplasmic fraction of Jurkat cells was mixed with the in vitro transcribed Y3 RNA and incubated for 2 h at 30 °C with or without activated caspase 3. Activated caspase 3 facilitated the cleavage of Y3 RNA and production of ASR.

### Activation of Y RNA cleavage is mediated by apoptotic stimulation

We previously observed elevation in ASR levels along with EBV reactivation when the ratio of dead cells was high. On the other hand, ASRs were expressed in not only EBV-positive cells but also EBV-negative cells. We hypothesized that Y RNA cleavage depends on cell death, which massively occurs during EBV lytic infection. Apoptotic cells were defined as Annexin V-positive cells (24). After 24 h of B cell receptor (BCR)-mediated lytic cycle induction, the number of apoptotic cells was increased in Akata cells (Annexin V [+] PI [−], 27.2%; Annexin V [+] PI [+], 25.8%; total Annexin V [+], 53.0%) as compared to cells without induction (Annexin V [+] PI [−], 5.6%; Annexin V [+] PI [+], 8.3%; total Annexin V [+], 13.9%) (Figure 1B). Apoptosis-associated Y RNA cleavage was verified by northern blotting (Figure 1C). Additionally, Y RNA cleavage was induced by activated caspase 3 in the cell-free system in vitro (Figure 1D). These results indicate that ASRs processing is activated in apoptotic cells and is coordinated by caspase3.

### Y RNA cleavage is independent of Drosha, Dicer, and RNase L

We previously reported that ASRs are processed independently from DROSHA (5). We examined whether ASRs were processed by DICER. To study the effect of DICER and DROSHA in processing, we performed siRNA-mediated knockdown of the genes and examined the expression of ASRs in apoptosis-inducing cells. No differences were observed in the processing of ASR between DICER-or DROSHA-knockdown or control cells (Figure 2A). Furthermore, the expression of ASR was unchanged between the wild-type and DICER-deficient MEF cells (Figure 2B). These results indicate that Y RNAs are cleaved by an unknown RNase but not DICER.

**Figure 2.**
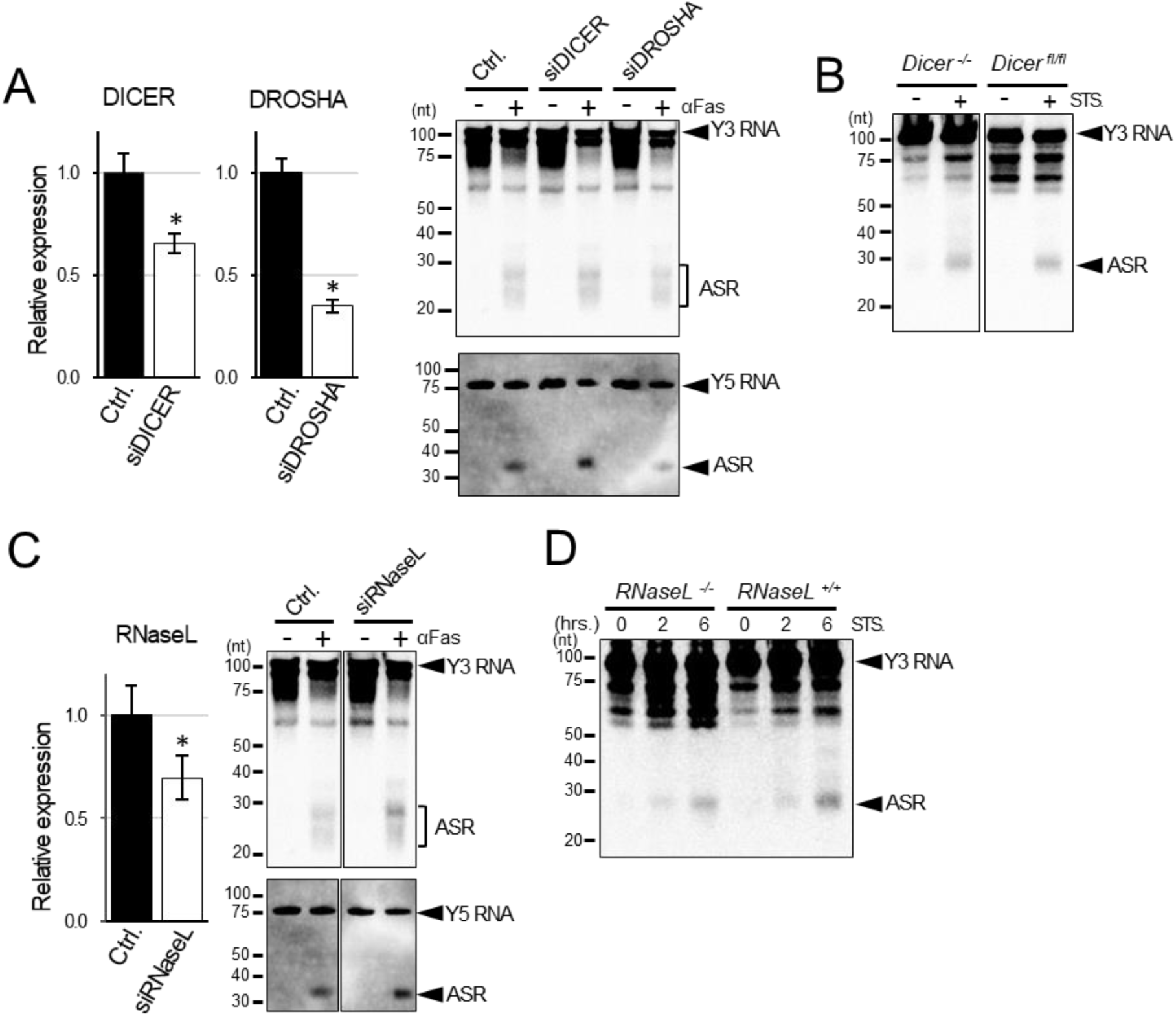
Y RNA processing is independent of DROSHA, DICER, and RNaseL. (A) Jurkat cells were transfected with control siRNA (Ctrl.), siDROSHA, or siDICER by NeonTM transfection system. Three days after transfection, the expression of target genes was examined by real-time PCR. The relative expression was normalized by GAPDH. Apoptosis was induced by 2.5 μg/mL anti-Fas antibody (clone: CH-11). After 6 h, total RNA was extracted and Y RNA fragments detected by northern blotting with an ASR probe. Upper panel, Y3 RNA and ASR; lower panel, Y5 RNA and ASR. (B) DICER-knockout or wild-type (floxed DICER) MEFs were cultivated with or without 10 µM staurosporine for 2 h, followed by the detection of Y3 RNA cleavage by northern blotting. (C) The expression of RNaseL decreased, as evaluated by real-time PCR, 3 days after siRNaseL transfection into Jurkat cells. Y3 RNA or Y5 RNA cleavage was also detected by northern blotting. (D) RNaseL‑knockout or wild-type MEFs were stimulated with 10 μM staurosporine. After 2 or 6 h, Y3 RNA cleavage was observed in both cells. For A and C, n = 3, * P < 0.05, two-tailed Student’s t-test; error bars, mean ± s.d.

Several viral single-strand RNAs are known to be cleaved by RNase L (25), which is activated during apoptosis (26). We examined whether ASRs were processed by RNase L. We performed siRNA-mediated knockdown of the genes and examined the expression of ASRs in apoptosis-inducing cells. No differences were observed in the processing of ASR between RNase L-knockdown and control cells (Figure 2C). Furthermore, ASR was detected in RNase L-knockout MEF cells following apoptotic induction by staurosporine (Figure 2D). Thus, ASR was processed in RNaseL-deficient condition, indicating that Y RNAs are processed by an unknown RNase but not RNase L.

### Identification of RNase and RNase inhibitor for Y RNA cleavage

The processing machinery of ASRs was investigated in a comprehensive manner. To identify the molecules responsible for Y RNA cleavage, the S-100 fraction from Jurkat cells was fractionated using saturated ammonium sulfate (SAS). Each fraction was treated with the activated caspase 3 and Y RNA processing examined. As shown in Figure 3A, a 2.3-fold increase in caspase 3-dependent Y RNA cleavage activity, calculated as ASR/Y3 RNA, was observed in the SAS 50-60% fraction as compared with that in the control without caspase 3 (1.03 versus 0.44) (Figure 3A). The SAS 50-60% fraction was separated by IEC and cleavage activities were examined with or without caspase 3. We failed to observe the dependency of caspase 3 for Y RNA processing in all fractions. The processing was examined in the reaction without caspase 3 (Figure 3B). Fractions e and f showed 6.36 and 6.84 processing activities, which were 2.56-and 2.75-fold higher, respectively, than the average in this assay (2.48). These fractions showed high RNase activity and were defined as RNase fraction (R.F.). From the results, it was hypothesized that molecules determining caspase 3 dependency comprised R.F. and other fractions, which may contain the inhibitory activity of RNase. To test this hypothesis, R.F. was mixed with other IEC fractions and incubated with the activated caspase 3 (Figure 3C). We detected 0.77, 0.36, 0.06, and 0.93 processing activities without caspase 3 with k to n fractions, which corresponded to 3.4, 7.4, 45.7, and 2.8-fold reduction, respectively, as compared to the average processing activity of R.F (2.6). Caspase 3-dependent processing was recovered upon mixing the fraction from k to m (Figure 3C). The processing activity of fraction m without and with caspase 3 was 0.06 and 1.52, respectively, indicating that a 26.4-fold increase in the processing activity was observed in the reaction with caspase 3 as compared to the reaction without caspase 3. These results suggest that RNase activity is inhibited by one or several factors from IEC fractions k to m and that the inhibition activity was deactivated by caspase 3.

**Figure 3.**
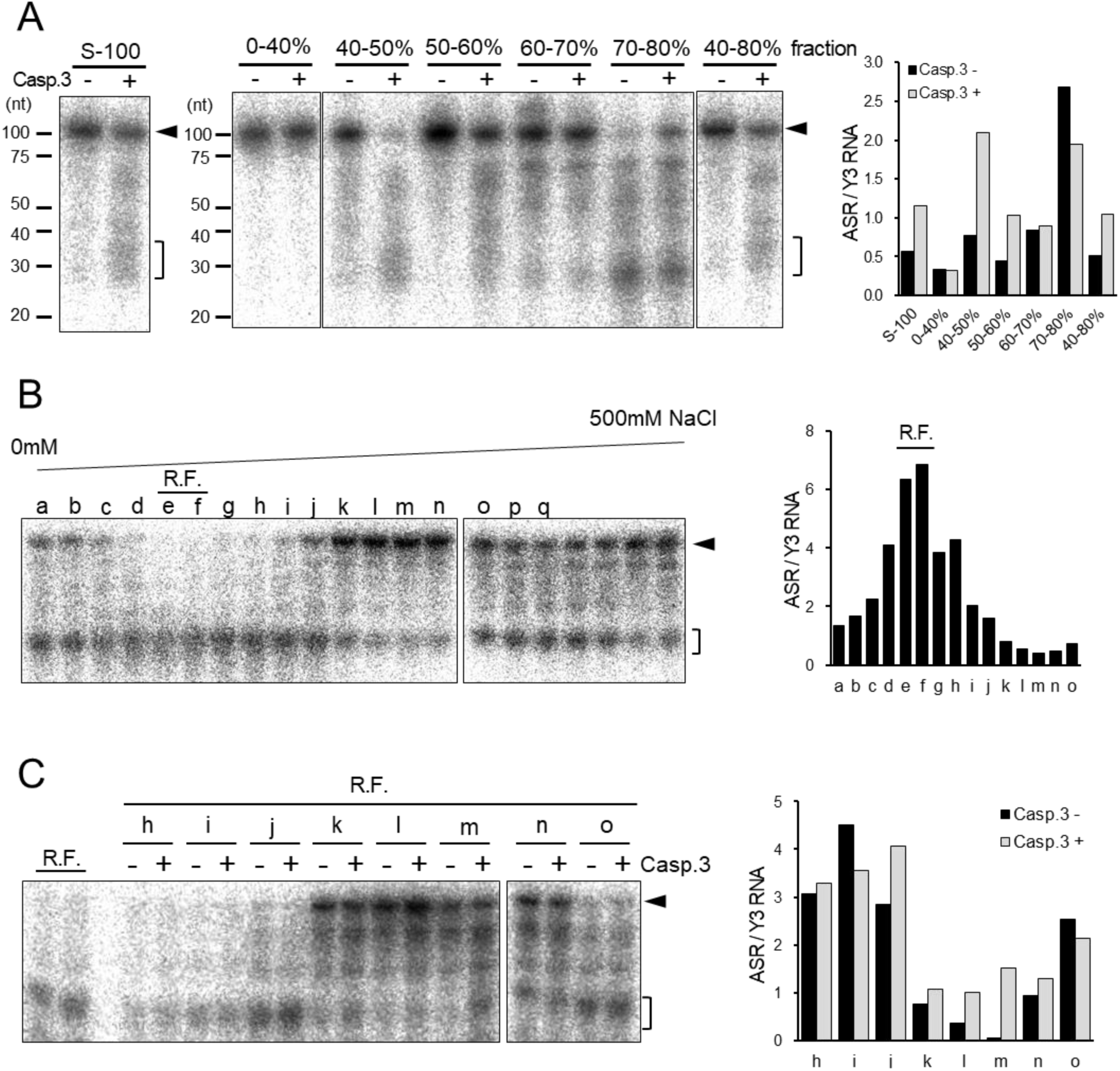
Y RNA processing involves ribonuclease and its inhibitor, which brings caspase 3 dependency for Y RNA processing. (A) The cytoplasmic fraction (S-100) of Jurkat cells was separated by ultracentrifugation and treated with 32P-labeled Y3 RNA for 2 h at 30°C with or without 0.25 unit of activated caspase 3. RNAs were purified from the reaction and separated by urea-PAGE (left). S-100 fraction was separated by ammonium sulfate fractionation. Caspase 3-dependent Y RNA processing was detected in 40-50%, 50-60%, and 40-80% fractions (middle). The ratio of ASR/Y3 RNA was calculated and plotted in a bar graph (right). (B) Ammonium sulfate fraction (50-60%) was dialyzed and injected into a strong anion-exchange chromatography column and eluted with a linear gradient of 0-500 mM NaCl. Equal volume of elution was analyzed and plotted on a bar graph. RNase activity was detected in fractions e and f, which were defined as RNase fraction (R.F.) (C) R.F. was combined and re-mixed with fractions from h to o and the caspase 3-dependent RNase activity was detected in fractions k, l, m, and n.

### PTBP1 inhibits Y RNA cleavage and caspase 3 deactivated the inhibition

It can be suggested that the inhibitor is truncated by caspase 3. Several proteins were shown to be bound to Y RNA (27). One of these proteins is PTBP1 (also known as hnRNP I), which binds to the loop region of Y RNA where it is cleaved during the biogenesis of ASRs (28). Furthermore, the cleavage of PTBP1 by caspase 3 was reported, but its significance has been undetermined (29). IEC fractions were analyzed by immunoblotting with anti-PTBP1 antibody and significant band densities were detected only in fractions with inhibitory activity (Figure 4A). To investigate the inhibitory activity of PTBP1 on Y RNA processing, recombinant PTBP1 protein which was extracted from 293T cells was incubated with the reaction solution containing Y RNA and R.F. fraction showing RNase activity. The Y3 RNA/ASR which indicates inhibitory activity of the cleavage, 2.8-fold was increased in the presence of PTBP1 but the inhibitory effected was cancelled by the activated caspase 3, indicating that Y RNA processing activities was inhibited by PTBP1 and caspase 3 deactivated this inhibition (Figure 4B). Moreover, the Y RNA cleavage was also examined with “uncleaved mutant PTBP1” which carrying amino acid substitution at caspase 3 target loci (D7A, D139A, and D172A) as described (29). While wildtype PTBP1 was cleaved by caspase3, the mutant PTBP1 was uncleaved in Figure 4C. In this condition, cleavage of Y3 RNA remained suppressed by the mutant PTBP1 with caspase 3, while that was proceeded by wildtype PTBP1 with caspase 3 (Figure 4D). These results demonstrated that the Y RNA processing inhibitor, PTBP1, is involved in apoptosis-dependent biogenesis of ASRs.

**Figure 4.**
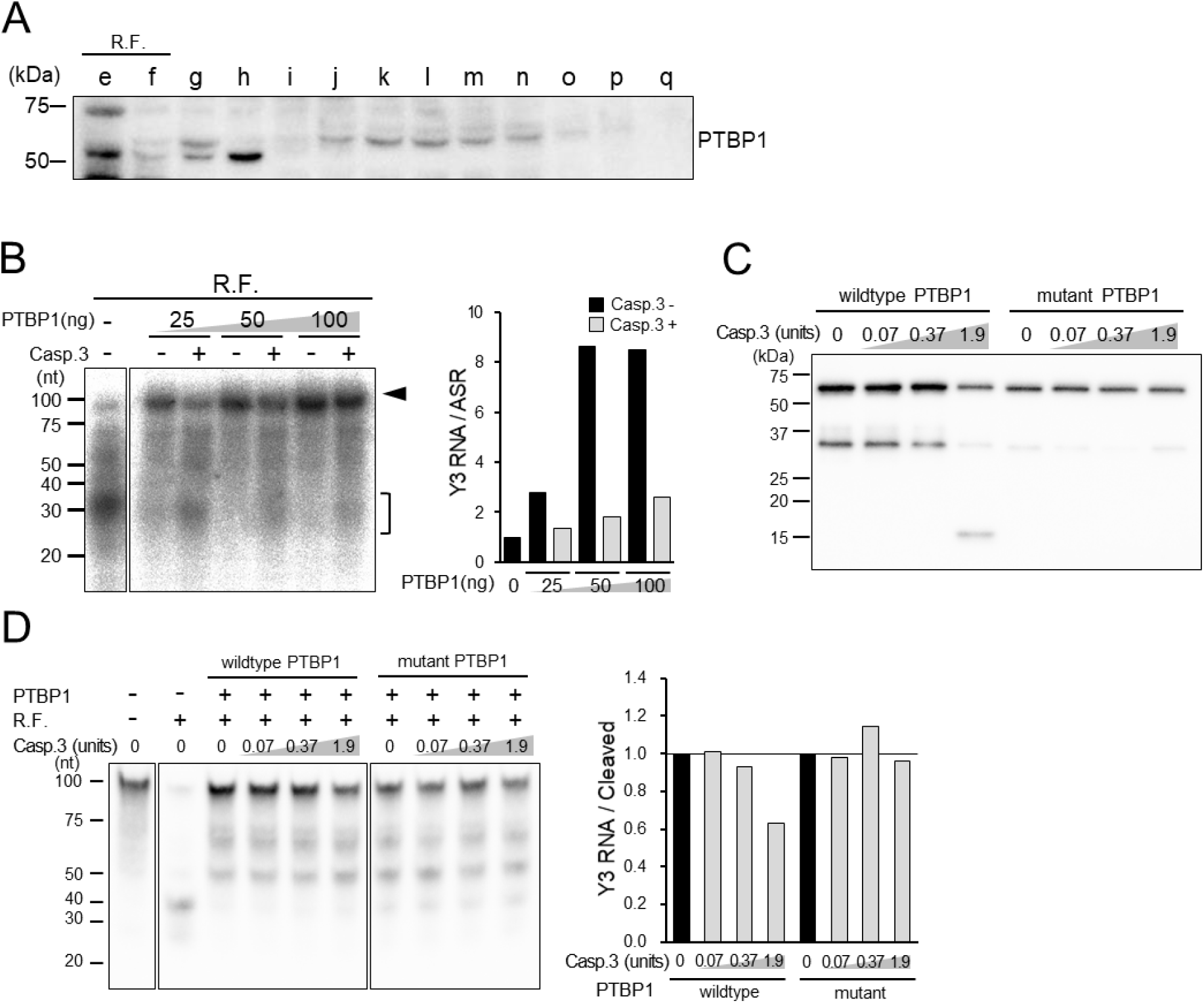
PTBP1 is responsible for the caspase 3-dependent Y RNA cleavage. (A) Equal volume of each IEC fraction was analyzed by western blotting with anti-PTBP1 antibody. (B) Recombinant PTBP1 was mixed with R.F., 32P-labeled Y3 RNA and 0.25 unit of activated caspase 3. After 2-h incubation at 30°C, Y3 RNA cleavage was detected by autoradiography (left). Y3 RNA / ASR ratios were calculated and normalized by the reaction without PTBP1 and plotted as a bar graph (right). (C,D) Wildtype or caspase 3 – resistant PTBP1, Y3 RNA, R.F. and activated caspase 3 were incubated for 2h at 37°C. Cleavage of PTBP1 was observed by Western blotting (C). Y3 RNA cleavage was detected by Northern blotting and cleavage activities were calculated as ratio of Y3 RNA / cleaved product, then normalized by the reaction without caspase 3, respectively (D).

## Discussion

Agotaxis small RNAs are a recently found class of small non-coding RNAs with the mechanism of action similar to that of miRNAs (5). ASRs were identified in EBV-infected cells and their expression was dramatically elevated during the lytic phase of EBV infection. According to recent next-generation sequencing studies, ASRs may be observed in various cells, indicating that ASRs processing may display some general functions not only in EBV infection but also in cellular biology. In this study, we found that the precursors of ASRs are non-coding Y RNAs, which are widely conserved and essential for the initiation of DNA replication. Y RNAs are known to be cleaved during apoptosis (9). The lytic phase of EBV infection is characterized by massive apoptosis. ASRs were thought to be generated during apoptosis from their precursor Y RNAs. Although the small RNA degradation or generation, which includes Y RNAs or U1 snRNA in apoptosis, was previously observed (7, 9), their cleavage enzymes, functions, and biological significance in apoptosis are unclear. In this study, we elucidated the cleaving machinery of Y RNA and processing machinery of ASR. As shown in Figure 1D, Y RNA cleavage may be induced from cytoplasmic extracts following treatment with caspase 3. The fractionation of the cell extract using SAS, IEC, and gel-filtration chromatography revealed at least two molecules, ribonuclease and PTBP1, that interact with the machinery. PTBP1 binds to the cleavage locus of Y RNAs and inhibits the access to RNase (Figure 5).

**Figure 5.**
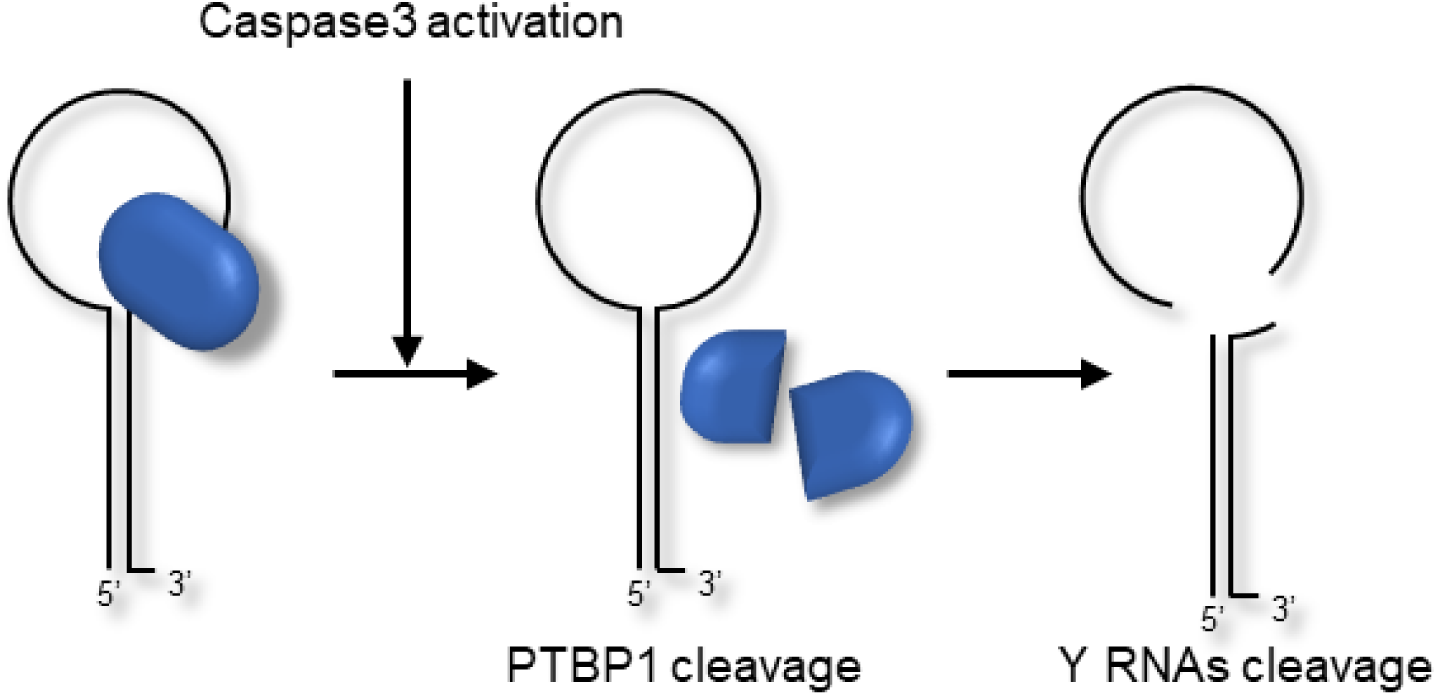
Schematic representation of Y RNA cleavage in apoptosis.

Despite similarities between the ASR and miRNA machineries for exerting their functions, canonical miRNA-processing enzymes for the biogenesis of miRNAs, DROSHA and DICER, had no role in the processing (Figure 2A,B). miRNAs are generally processed by DICER, which interacts with AGO protein when miRNA loads onto RNA-induced silencing complex (RISC). We have previously shown that the most distinctive feature of ASRs was to be selectively loaded onto AGO1. It would be interesting to know whether the molecules involved in the processing of ASRs, including unknown RNases, are related to selective AGO1 loading.

We performed mass spectrometry analysis to determine RNase using samples from gel-filtration chromatography and IEC R.F (Figure 3B, data not shown). However, we failed to identify the molecules, owing to the detection of excess of protein species. To overcome this problem, additional fractionation or other screening methods would be needed. Chromatin fragmentation is a well characterized event in apoptosis. Caspase-activated DNase (CAD)/DNA fragmentation factor beta (DFFB) is an endonuclease specifically activated in apoptosis by caspase 3 (30); and CAD lacks a caspase target site. On the other hand, the CAD inhibitor ICAD displays caspase 3 target site and usually binds to CAD to prevent its DNase activity (3,31) in a manner similar to Y RNA cleavage machinery, because the apoptotic cleavage of Y RNAs showed that the inhibitor PTBP1 is cleaved by caspase 3.

Experiments using knockout mice revealed that the systemic deficiency of DNase II causes DNA accumulation in macrophages, resulting in excess innate-immune response (32). In addition, systemic autoimmune diseases such as Sjogren’s syndrome and systemic lupus erythematosus are characterized by autoantibodies against nucleic acids or ribonucleocomplexes in the patient’s serum. Ro60 and La (also known as SSA/SSB) which autoantibodies are detected in the serum of almost every Sjogren’s syndrome patient are Y RNA-binding proteins. Although the relationship between the autoantigen and pathogenesis is poorly understood, Y RNA cleavage could have some roles to suppress the exposure of autoantigens to immunosurveillance (33). To examine whether Y RNA cleavage in apoptosis is involved in the generation of autoantigens or pathogenesis of autoimmune diseases, the establishment of knock-in mice that express caspase 3-resistant PTBP1, is now in progress.

In summary, we investigated the molecular machinery of apoptotic Y RNA cleavage. Y RNAs are cleaved by a cytoplasmic endoribonuclease and its inhibitor PTBP1 induces apoptotic Y RNA cleavage via its truncation by caspase 3.

## Experimental procedures

### Cells

Epstein-Barr virus-positive Akata cells and Jurkat cells were maintained in RPMI-1640 medium (Wako) supplemented with 10% (v/v) fetal bovine serum (FBS), 50 U/mL penicillin, and 50 μg/mL streptomycin. DICER-or RNaseL (kindly gifted by Dr. R. Silverman) -deficient mouse embryonic fibroblasts (MEFs) and HEK293T cells were maintained in DMEM (Nacalai Tesque) supplemented with 10% (v/v) FBS, 50 U/mL penicillin, and 50 μg/mL streptomycin.

### Apoptosis and lytic phase induction

To induce apoptosis, MEF cells were treated with 10 μM staurosporine (Wako) at 37°C for 2-6 h. To induce the EBV lytic phase, Akata cells were stimulated with rabbit anti-human IgG polyclonal antibody (20 mg/mL) (Dako) at 37°C for 24 h. Apoptosis was detected by propidium iodide (PI)/allophycocyanin (APC)-Annexin V (Merck and BD Bioscience) staining and analyzed by a flow cytometer (FACSVerse, BD Bioscience).

### Northern blotting

The purified total RNA (5 μg) was mixed with an equal volume of gel loading buffer II (Ambion) and denatured at 95°C for 5 min. Denatured samples were separated by electrophoresis using 15% urea gel and transferred onto Hybond N+ membrane (GE Healthcare) in cold room. Hybridization was performed with ULTRAhyb™ buffer (Ambion) at 37°C for overnight. Digoxigenin (DIG)-labeled Locked Nucleic Acid (LNA) probe for ASR derived from Y3 RNA was purchased from EXIQON. Hybridized probes were detected by anti-DIG-AP, Fab fragments, and CSPD substrate (Roche). The detection probe for ASR derived from Y5 RNA was prepared using the DIG oligonucleotide tailing kit, 2nd generation (Roche), according to the manufacturer’s instructions.

### Transfection with siRNAs

We purchased siRNAs targeting human DROSHA and DICER from OriGene Technology, Inc. siRNaseL, was obtained from Hokkaido System Science. AccuTarget^TM^ Negative Control siRNA (Bioneer) was used as control. Transfection was performed with the Neon transfection system (Invitrogen) according to the manufacturer’s protocol. Target genes were evaluated by real-time PCR. Sequences of specific siRNAs are described in Supplemental Table S1.

### Quantification of gene expression

Total RNA was purified from cells, using Sepasol-RNA I Super G (Nacalai Tesque), and reverse transcribed using the High Capacity cDNA transcription kit (Thermo Fisher Scientific). Real-time PCR was performed using THUNDERBIRD SYBR qPCR Mix (TOYOBO) with StepOnePlus real-time PCR system (Applied Biosystems). Threshold cycle (CT) values were calibrated to β-actin and analyzed by the 2^−^⊿⊿^CT^ method. Gene-specific primers are described in Supplemental Table S1.

### Protein fractionation and ribonuclease activity assay

Jurkat cells were suspended in an equal volume of a buffer containing 50 mM PIPES-KOH (pH 7.4), 50 mM KCl, 5 mM EGTA, 2 mM MgCl_2_, 1 mM DTT, 20 µM cytochalasin B (Wako), and protease inhibitor cocktail (Merck). The suspension was immediately frozen in ice-cold iso-propanol bath and melted on ice. The thawed sample was disrupted by Dounce homogenizer (KONTES) with 50 strokes. The freeze, thaw, and disrupt cycle was repeated. The lysate was centrifuged at 10,000 ×*g* for 12 min at 4°C and moved to a new tube. The supernatant was separated by ultracentrifugation at 100,000 ×*g* and 4°C for 90 min. The collected supernatant was subjected to ion-exchange chromatography (IEC) with HiTrap Q column (GE healthcare) and eluted by a linear gradient of 0-500 mM NaCl. ^32^P-labeled Y3 RNA was prepared using an in vitro transcription T7 kit (TaKaRa). Each fraction and ^32^P-labeled Y3 RNA were mixed in 10 mM HEPES-KOH pH7.0, 50 mM NaCl, 2 mM MgCl_2_, 20% glycerol, 40 mM beta-glycerophosphate, 5 mM DTT, and 1 mg/mL bovine serum albumin (BSA) and incubated with or without caspase 3 (kindly gifted by Dr. S. Nagata) for 2 h at 30°C. RNA was purified from the reactions by acid-phenol and ethanol precipitation and separated by 15% acrylamide-denatured gel (urea). The gel was exposed to imaging plate and analyzed by FLA-2000 (FUJIFILM).

### Western blotting

Each sample was separated by sodium dodecyl sulfate polyacrylamide gel electrophoresis (SDS-PAGE). The separated proteins were transferred onto a polyvinylidene difluoride (PVDF) membrane (Merck) and the membrane was blocked using PVDF blocking reagent for Can Get Signal (TOYOBO) at 25°C for 1 h. Blotting was performed with anti-PTBP1 antibody (RN011P, MBL). Horseradish peroxidase (HRP)-conjugated secondary antibody or streptavidin was added to the membranes and the peroxidase activity detected using Immobilon western chemiluminescent HRP substrate (Merck).

### Recombinant PTBP1 preparation

We cloned PTBP1 into pcDNA3-Flag and transfected the vector into 2 × 10^7^ HEK293T cells, using polyethylenimine “MAX” (Polysciences). After 2 days of transfection, nuclear lysate was mixed with anti-FLAG M2 affinity gel (Merck) and eluted by incubation with 3× Flag peptide (Merck). A cDNA of StrepTagII-tagged PTBP1 was cloned into pET49 plasmid. To generate the vector expressing caspase 3-resistant PTBP1, site-directed mutagenesis was performed using Pfu Turbo DNA Polymerase (Agilent Technologies), with three primer pairs carrying the nucleotides for D7A, D139A, and D172A mutations. These vectors were transformed into *Escherichia coli* BL21(DE3) strain. The lysates were injected into StrepTrap HP column (GE Healthcare) and eluted by 2.5 mM dethtiobiotin (Merck).

## Acknowledgements

The authors would like to thank Dr Shigekazu Nagata, Dr Nobuyuki Miyasaka, Dr. Robert H. Silverman, Dr Jun Suzuki, Ms. Masako Takamatsu, Ms. Natsumi Kurosaki, and Educational Research Center of Tokai University School of Medicine for their technical assistance. This work was supported by PRESTO, AMED-PRIME, and the Research Program on Hepatitis from Japan Agency for Medical Research and Development, AMED to A.K. and assignments.

## conflict of interest

The authors declare that they have no conflicts of interest with the contents of this article.

## Author contributions

JO. and AK designed research and wrote the article. JO, YS, MT, HM, MO MO and TI performed experiments. Bioinformatics analysis was performed by AK The manuscript was reviewed by all authors.

## FOOTNOTES

This work was supported by JSPS KAKENHI (17H04212 to A.K.), Japan Agency for Medical Research and Development – Precursory Research for Innovative MEdical care (15665962 to A.K.) and the Research Program on Hepatitis from Japan Agency for Medical Research and Development (16768629 to M.O.). Funding for open access charge: JSPS KAKENHI.

The abbreviations used are: PTBP1, Polypyrimidine tract-binding protein 1; AGO, Argonaute; rRNA, ribosomal RNA; piRNA, piwi-interacting RNA

## References

1. Jacobson, M. D., Weil, M., and Raff, M. C. (1997) Programmed cell death in animal development. Cell 88, 347–354

2. Ellis, H. M., and Horvitz, H. R. (1986) Genetic control of programmed cell death in the nematode C. elegans. Cell 44, 817–829

3. Enari, M., Sakahira, H., Yokoyama, H., Okawa, K., Iwamatsu, A., and Nagata, S. (1998) A caspase-activated DNase that degrades DNA during apoptosis, and its inhibitor ICAD. Nature 391, 43–50

4. Kerr, J. F., Wyllie, A. H., and Currie, A. R. (1972) Apoptosis: a basic biological phenomenon with wide-ranging implications in tissue kinetics. Br J Cancer 26, 239–257

5. Degen, W. G., Pruijn, G. J., Raats, J. M., and van Venrooij, W. J. (2000) Caspase-dependent cleavage of nucleic acids. Cell Death Differ 7, 616–627

6. Houge, G., Døskeland, S. O., Bøe, R., and Lanotte, M. (1993) Selective cleavage of 28S rRNA variable regions V3 and V13 in myeloid leukemia cell apoptosis. FEBS Lett 315, 16–20

7. Degen, W. G., Aarssen, Y., Pruijn, G. J., Utz, P. J., and van Venrooij, W. J. (2000) The fate of U1 snRNP during anti-Fas induced apoptosis: specific cleavage of the U1 snRNA molecule. Cell Death Differ 7, 70–79

8. Casciola-Rosen, L. A., Miller, D. K., Anhalt, G. J., and Rosen, A. (1994) Specific cleavage of the 70-kDa protein component of the U1 small nuclear ribonucleoprotein is a characteristic biochemical feature of apoptotic cell death. J Biol Chem 269, 30757–30760

9. Rutjes, S. A., van der Heijden, A., Utz, P. J., van Venrooij, W. J., and Pruijn, G. J. (1999) Rapid nucleolytic degradation of the small cytoplasmic Y RNAs during apoptosis. J Biol Chem 274, 24799–24807

10. Kowalski, M. P., and Krude, T. (2015) Functional roles of non-coding Y RNAs. Int J Biochem Cell Biol 66, 20–29

11. Christov, C. P., Gardiner, T. J., Szüts, D., and Krude, T. (2006) Functional requirement of noncoding Y RNAs for human chromosomal DNA replication. Mol Cell Biol 26, 6993–7004

12. Krude, T., Christov, C. P., Hyrien, O., and Marheineke, K. (2009) Y RNA functions at the initiation step of mammalian chromosomal DNA replication. J Cell Sci 122, 2836–2845

13. Sim, S., Weinberg, D. E., Fuchs, G., Choi, K., Chung, J., and Wolin, S. L. (2009) The subcellular distribution of an RNA quality control protein, the Ro autoantigen, is regulated by noncoding Y RNA binding. Mol Biol Cell 20, 1555–1564

14. Schur, P. H., Moroz, L. A., and Kunkel, H. G. (1967) Precipitating antibodies to ribosomes in the serum of patients with systemic lupus erythematosus. Immunochemistry

15. Uchiumi, T., Traut, R. R., Elkon, K., and Kominami, R. (1991) A human autoantibody specific for a unique conserved region of 28 S ribosomal RNA inhibits the interaction of elongation factors 1 alpha and 2 with ribosomes. J Biol Chem 266, 2054–2062

16. Lerner, M. R., and Steitz, J. A. (1979) Antibodies to small nuclear RNAs complexed with proteins are produced by patients with systemic lupus erythematosus. Proc Natl Acad Sci U S A 76, 5495–5499

17. Clark, G., Reichlin, M., and Tomasi, T. B. (1969) Characterization of a soluble cytoplasmic antigen reactive with sera from patients with systemic lupus erythmatosus. J Immunol 102, 117–122

18. Bartel, D. P. (2004) MicroRNAs: genomics, biogenesis, mechanism, and function. Cell 116, 281–297

19. Yang, N., and Kazazian, H. H. (2006) L1 retrotransposition is suppressed by endogenously encoded small interfering RNAs in human cultured cells. Nat Struct Mol Biol 13, 763–771

20. Ku, H. Y., and Lin, H. (2014) PIWI proteins and their interactors in piRNA biogenesis, germline development and gene expression. Natl Sci Rev 1, 205–218

21. Yamakawa, N., Okuyama, K., Ogata, J., Kanai, A., Helwak, A., Takamatsu, M., Imadome, K., Takakura, K., Chanda, B., Kurosaki, N., Yamamoto, H., Ando, K., Matsui, H., Inaba, T., and Kotani, A. (2014) Novel functional small RNAs are selectively loaded onto mammalian Ago1. Nucleic Acids Res 42, 5289–5301

22. Mulligan, G. J., Guo, W., Wormsley, S., and Helfman, D. M. (1992) Polypyrimidine tract binding protein interacts with sequences involved in alternative splicing of beta-tropomyosin pre-mRNA. J Biol Chem 267, 25480–25487

23. van Gelder, C. W., Thijssen, J. P., Klaassen, E. C., Sturchler, C., Krol, A., van Venrooij, W. J., and Pruijn, G. J. (1994) Common structural features of the Ro RNP associated hY1 and hY5 RNAs. Nucleic Acids Res 22, 2498–2506

24. van Engeland, M., Kuijpers, H. J., Ramaekers, F. C., Reutelingsperger, C. P., and Schutte, B. (1997) Plasma membrane alterations and cytoskeletal changes in apoptosis. Exp Cell Res 235, 421–430

25. Zhou, A., Paranjape, J., Brown, T. L., Nie, H., Naik, S., Dong, B., Chang, A., Trapp, B., Fairchild, R., Colmenares, C., and Silverman, R. H. (1997) Interferon action and apoptosis are defective in mice devoid of 2′,5′-oligoadenylate-dependent RNase L. EMBO J 16, 6355–6363

26. Castelli, J. C., Hassel, B. A., Maran, A., Paranjape, J., Hewitt, J. A., Li, X. L., Hsu, Y. T., Silverman, R. H., and Youle, R. J. (1998) The role of 2′-5′ oligoadenylate-activated ribonuclease L in apoptosis. Cell Death Differ 5, 313–320

27. Köhn, M., Pazaitis, N., and Hüttelmaier, S. (2013) Why YRNAs? About Versatile RNAs and Their Functions. Biomolecules 3, 143–156

28. Fabini, G., Raijmakers, R., Hayer, S., Fouraux, M. A., Pruijn, G. J., and Steiner, G. (2001) The heterogeneous nuclear ribonucleoproteins I and K interact with a subset of the ro ribonucleoprotein-associated Y RNAs in vitro and in vivo. J Biol Chem 276, 20711–20718

29. Back, S. H., Shin, S., and Jang, S. K. (2002) Polypyrimidine tract-binding proteins are cleaved by caspase-3 during apoptosis. J Biol Chem 277, 27200–27209

30. Nicholson, D. W., Ali, A., Thornberry, N. A., Vaillancourt, J. P., Ding, C. K., Gallant, M., Gareau, Y., Griffin, P. R., Labelle, M., and Lazebnik, Y. A. (1995) Identification and inhibition of the ICE/CED-3 protease necessary for mammalian apoptosis. Nature 376, 37–43

31. Sakahira, H., Enari, M., and Nagata, S. (1998) Cleavage of CAD inhibitor in CAD activation and DNA degradation during apoptosis. Nature 391, 96–99

32. Kawane, K., Fukuyama, H., Yoshida, H., Nagase, H., Ohsawa, Y., Uchiyama, Y., Okada, K., Iida, T., and Nagata, S. (2003) Impaired thymic development in mouse embryos deficient in apoptotic DNA degradation. Nat Immunol 4, 138–144

33. Reed, J. H., Sim, S., Wolin, S. L., Clancy, R. M., and Buyon, J. P. (2013) Ro60 requires Y3 RNA for cell surface exposure and inflammation associated with cardiac manifestations of neonatal lupus. J Immunol 191, 110–116

